# Evidence for an ancient master sex determination gene in Hymenoptera

**DOI:** 10.64898/2025.12.24.696408

**Authors:** Sean M. Velasquez, Isaac J. Walker, Tyler Allen McClure, Leia Isabela S. Pineda, Maxence Gérard, Virginia V. Russell, Paige Camaya, Gwenndolyn D. J. Campbell, Nicholas V. Fernandes, Anaya Flowers, Ashley N. Raymond, Sam S. Rico, Anton Taylor, Ramona C. Velazquez, Alexander Kramer, Christopher J. Condon, Russell Corbett-Detig, Alan Brelsford, Noelle Anderson, Scott William Roy

## Abstract

The vast majority of Hymenopterans determine sex by haplodiploidy, in which males and females develop from haploid unfertilized and diploid fertilized eggs, respectively. At the molecular level, the majority of Hymenoptera are thought to determine sex by so-called Complementary Sex Determination (CSD). Under CSD, sex is determined by the feminizing effects of one or more allele-rich loci, in which the feminizing function is imparted by the interactions of functionally distinct alleles. Because of high allelic diversity, most diploids are heterozygous and develop as females, whereas haploid individuals develop as males, as do rare homozygous diploids. Despite diverse empirical results suggesting that CSD is widespread in Hymenopterans, and substantial theoretical and empirical study of the consequences of CSD, the molecular biology of CSD has received little attention outside of the honeybee, *Apis mellifera*. Available evidence provides support for divergent perspectives ranging from a single (nearly) universally-conserved CSD locus or a pattern dominated by evolutionary turnover with different genes serving as the CSD gene(s) in different lineages. Recently, a novel non-coding RNA gene named ANTSR was shown to serve as a CSD gene in different ant lineages, and the authors suggested that ANTSR might serve as a more broadly-conserved CSD gene. To assess the role of ANTSR role in CSD across diverse Hymenoptera, we studied syntenic conservation and polymorphism patterns in the broad ANTSR locus, genome-wide sequence from available male and female diploid *Formica* ants, and data from a sibling cross for *Bombus terrestris*. We find evidence that ANTSR is a conserved CSD gene across diverse Aculeata, the largest group of Hymenoptera, and provide preliminary evidence that *Formica* may have multiple CSD loci including ANTSR.

## INTRODUCTION

The insect group Hymenoptera is an extremely ecologically important group of insects, including bees, ants, wasps, sawflies and their allies. It includes many solitary species, as well as nearly all instances of highly social insects. Almost all Hymenoptera determine sex by haplodiploidy, in which males and females are haploid and diploid, respectively, due to developing from unfertilized and fertilized eggs (Heimpel et al. 2008).

Despite sharing the haplodiploidy sex determination mechanism, different Hymenopterans use different mechanisms for sex determination at the molecular level (Asplen et al. 2009). The best studied mode of sex determination in Hymenopterans is so-called Complementary Sex Determination (CSD) (Whiting 1943). In the simplest case, female development depends on heterozygosity at a single allele-rich locus, with heterozygous diploid individuals developing as females while haploid and homozygous diploid individuals developing as males. In contrast to this “single-locus” CSD case, multi-locus CSD is also known, in which heterozygosity at at least one of multiple feminizing loci is sufficient to trigger feminization (Asplen et al. 2009; Matthey-Doret et al. 2019; Miyakawa et al. 2018). Most, but not all, haplodiploid Hymenopterans are thought to use CSD (Asplen et al. 2009). The best characterized exception is found in Nasonia wasps, in which sex is determined due to a feminizing gene that is genomically imprinted, being expressed only from the paternally-inherited chromosome (Verhulst et al. 2010); thus, fertilized eggs contain an expressed allele of this feminizing gene, whereas unfertilized eggs contain only a silenced copy and thus develop as males. This gene was recently mapped, and found to be a recent duplicate from a p53-related gene not previously known to be involved in sex determination (Zou et al. 2020).

While organismal evidence, in particular the observation of diploid males, suggests that CSD is widespread across Hymenoptera, the specific CSD gene(s) has only been mapped in a small number of species (Matthey-Doret et al. 2019; Miyakawa et al. 2015; Beye et al. 2003; Otte et al. 2023; Rönneburg et al. 2025). First, the CSD gene was mapped in the honeybee *A. mellifera*, and was found to be a duplicate copy of the insect-wide female sex determination gene *Fem*, and named *CSD* (Beye et al. 2003). Mapping of a CSD in the ant *Vollenhovia emeryi* found two CSD loci, one of which also corresponded to a duplicate of *Fem*, and a second of which showed no obvious candidate sex determination gene (Miyakawa et al. 2015). The finding of homologous *Fem* genes in very divergent lineages (ants versus bees) performing the same function raised the possibility that Hymenoptera could largely share an ancestral, conserved master sex determination gene (Miyakawa et al. 2015), as found in some other animal lineages including mammals (Wallis et al. 2008), and in contrast to other lineages which show rapid turnover of sex determination mechanism (Mank and Avise 2009).

However, the notion of an ancestral *Fem-*related CSD locus has been complicated by further studies of the second CSD locus found in *V. emeryi*. This same general genomic locus was subsequently found to also serve as a CSD gene in three other ant lineages (Huang et al. 2014; Pan et al. 2024; Lacy et al. 2025), with a recent paper going so far as to functionally characterize the sex determination gene itself (Pan et al. 2024). This apparent CSD gene, named Ant Noncoding Transformer Regulator, or ANTSR, was found to be a lncRNA gene, with no known homology to other sex-relevant genes. Expression of ANTSR affects splicing of the core sex determination genes, indicating that ANTSR is a modulator of the core sex determination gene expression pattern (Pan et al. 2024). While this finding is consistent with ANTSR serving as an ancestral master CSD gene (as suggested by Pan et al. 2024), it did not exclude that, like the *Fem* gene family or *Wom*, ANTSR was only secondarily co-opted as a master sex determination gene.

In this study, we sought to test whether ANTSR might be an ancient, conserved sex determination gene across Hymenoptera. Using sibling bumblebee crosses and nucleotide diversity profiles across diverse Hymenoptera, we provide evidence that ANTSR is an ancestral CSD sex determination gene broadly conserved within the Hymenoptera group Aculeata.

## RESULTS

### Conservation of synteny at the ANTSR locus across Aculeata

Previous work showed conservation of a block of eight syntenic protein-coding genes flanking the ANTSR locus (Pan et al. 2024). This finding is interesting in its own light as preliminary data suggests that this represents an elevated degree of syntenic conservation (data not shown) and practically useful for identifying the ANTSR locus, given fast rates of sequence evolution of lncRNAs (e.g., Schüler et al. 2014). We analyzed genome assemblies and annotation files from 112 Hymenopteran species retrieved from the NCBI database. For most species within Aculeata (ants, bees, vespoid wasps, and their allies), annotated gene models confirmed conservation of microsynteny of the genes flanking ANTSR (Figure 1; Supplemental Table 1). Notably, given the reliance of this analysis on some low-quality genome assemblies, it remains possible that some of the negative findings are false negatives. Within Aculeata, the order and orientation of flanking genes were generally maintained, supporting broad conservation of the ANTSR genomic context. In a minority of Aculeata species, the flanking genes were positioned at the ends of separate contigs, preventing confirmation or rejection of gene order due to assembly limitations. Outside Aculeata, no clear conservation of the syntenic block was found. Although orthologous genes were sometimes found in proximity, the canonical syntenic structure observed in Aculeata genomes was absent. Together, these findings indicate that the ANTSR locus occupies a conserved genomic position across Aculeata, while the local genomic context appears to have diverged in other Hymenopteran groups (Figure 1; Supplemental Table 1).

**Figure 1.**
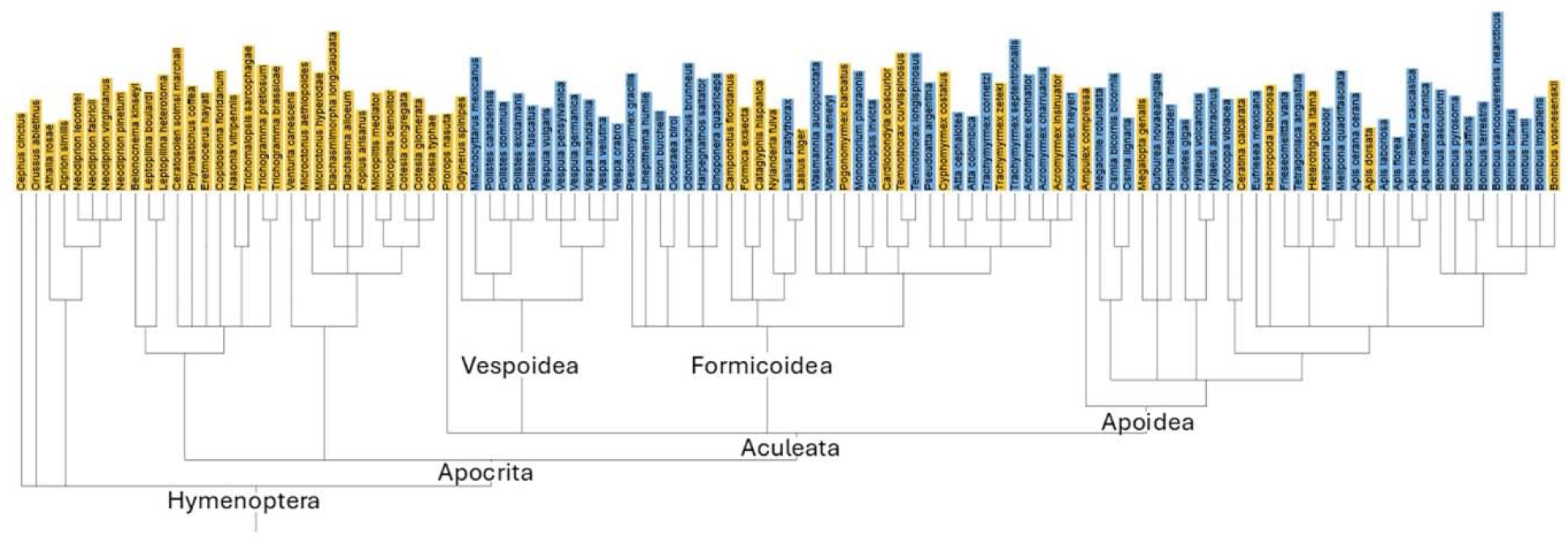
Diverse species within Aculeata show conserved synteny surrounding the ANTSR genomic region. Blue and yellow bars indicate presence and absence of synteny, respectively, across 112 Hymenopteran taxa. See Supplemental Table 1 for details.

### Elevated nucleotide diversity at the ANTSR locus across Aculeata

Selection against low-fitness diploid males is expected to drive balancing selection at CSD loci. Balancing selection is generally associated with increased nucleotide diversity, due both to functional diversification and to increased allelic age. Thus, we tested whether ANTSR showed elevated nucleotide diversity in Aculeata (as previously found in *L. humile* by Pan et al. 2024).

To evaluate nucleotide diversity at the ANTSR locus, whole-genome resequencing data from 37 species were mapped to their respective reference genomes and analyzed for nucleotide polymorphism (Figure 2, Supplemental Figure 1). Read mapping and polymorphism analysis revealed elevated nucleotide polymorphism near the ANTSR locus in 20 species, spanning the three major Aculeata superfamilies: Formicoidea, Apoidea and Vespoidea. In several additional species, signals of elevated polymorphism signals were detectable but weaker or partially obscured by limited coverage or assembly quality. These cases were classified as inconclusive rather than negative, as technical limitations could not be distinguished from biological absence of elevated polymorphism.

**Figure 2.**
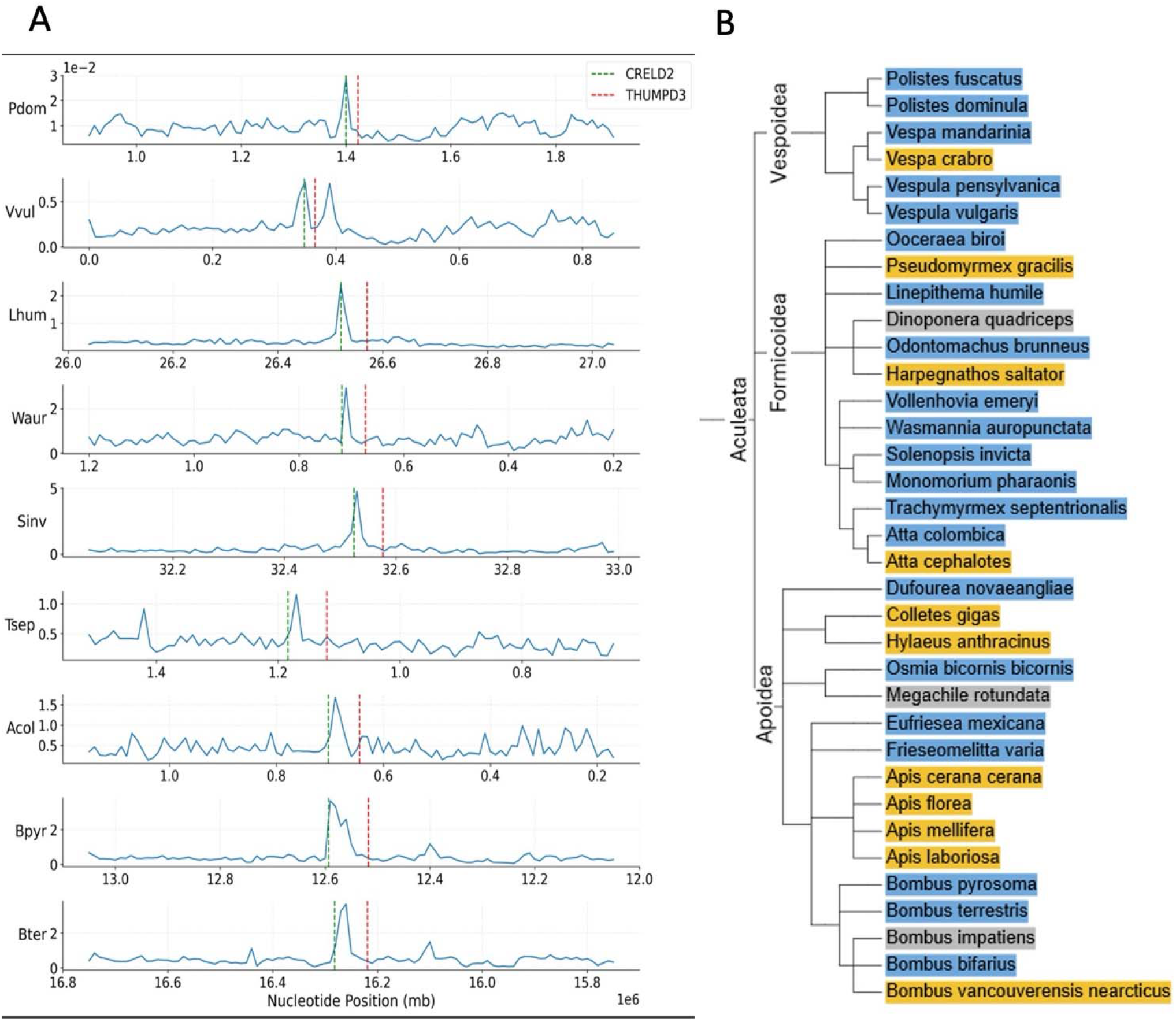
Diverse species within Aculeata show elevated nucleotide intraspecific diversity near the ANTSR locus. A. Selected examples of elevated polymorphism at the ANTSR locus across ants, bees and wasps. The dotted lines delineate the region between the neighboring protein-coding genes CRELD2 and THUMPD3. Estimated pairwise divergence between the reference genome and a second available isolate are shown. Species are the vespoid wasps *Polistes dominula* and *Vespula vulgaris*, the ants *Linepithema humile, Solenopsis invicta, Wasmannia auropunctata, Atta colombica* and *Trachymrmex septentrionalis*, and the bees *Bombus pyrosoma* and *Bombus terrestris*. B. Elevated polymorphism is found at or near the ANTSR locus across diverse Aculeata species (blue), though species without elevated polymorphism (yellow) or with ambiguous results (grey) are also observed.

### Evaluating turnover of ANTSR as the CSD locus

We also sought to study evolutionary turnover of the CSD locus, as seen in the honeybee *Apis mellifera*, in which a duplicate of *Fem*, and not ANTSR, serves as the CSD gene (Beye et al. 2002). Indeed, no signature of elevated polymorphism at the ANTSR locus is observed in *Apis mellifera* or several relatives (Figure 2). While several individual species of Aculeata also lack a clear signature of elevated polymorphism at the ANTSR locus (Figure 2), we found no additional clear cases of a clade of species lacking elevated ANTSR polymorphism. While these individual species are candidates for additional cases of CSD turnover, caution is appropriate, since several other possibilities exist, including problems with poorly-annotated datasets; chance allele sharing between the resequencing sample and the reference genome; newly-arising alleles with minimal nucleotide divergence; and recombination leading to homogenization between sites not directly involved in the CSD mechanism.

### Elevated polymorphism in diploid females from a sibling bumblebee cross

We generated diploid male and diploid female offspring from crossing two different pairs of siblings of the bumblebee *Bombus terrestris*, and confirmed ploidy by flow cytometry. We obtained 14 diploid males and 9 workers for one family, and 4 diploid males and a single worker for a second family. We conducted Illumina DNA sequencing of the single female from the small family and for pools of each set of diploid males and females (3 total pools). For each family, we identified polymorphic SNPs segregating within the daughter(s), and estimated the minor allele frequency in their diploid brothers, and averaged these values over 1 Mb windows. For the larger family (Figure 3A), the lowest estimated frequency falls at the ANTSR locus (red vertical line on Chr 2). Similarly, for the small family (Figure 3B), the ANTSR locus shows very low minor allele frequences, though other regions show comparably low minor allele frequencies, likely due to random fluctuations due to the small number of males.

**Figure 3.**
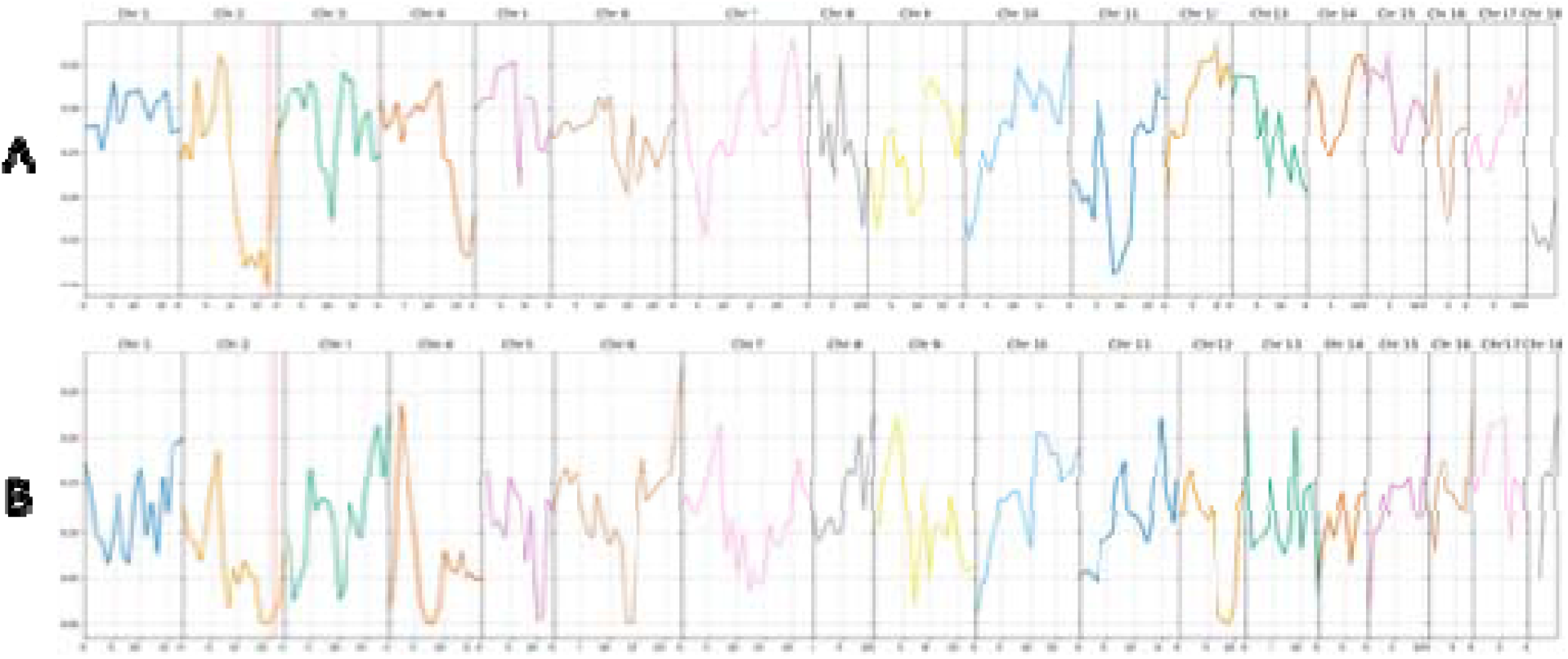
Estimated minor allele frequencies in diploid males, at polymorphic sites in their diploid sisters. A. The large family (14 males, 9 females). B. The small family (4 males, 1 female). The red vertical line represents the ANTSR locus.

### Naturally occurring diploid males and their nestmates in Formica cinerea suggest multiple CSD loci including ANTSR

We used available population RAD-seq data from the ant *Formica cinerea* to test the likelihood that ANTSR served as a CSD locus in that species. Analysis of heterozygosity at confident SNP sites identified four putative diploid males. For 774 females and the 4 diploid males, we assessed the distance between the nearest upstream and downstream heterozygous SNPs for each individual, which is equivalent to the maximum possible run of heterozygosity near ANTSR (given that a true assessment of ROH is not possible because of the sparsity of RAD-seq markers). All 4 diploid males fell within the top 6/778 individuals for this metric (*p* < 0.001 by a Mann-Whitney U test), consistent with homozygosity at ANTSRs leading to male development, and thus with a role for ANTSR in CSD (Figure 4). Interestingly, 3/4 diploid males derived from the same colony (ARN2102); 3/9 females from the same colony fell within the top 20 values out of 774 females (*p* < 0.01), though there was still a signal for males to have higher values than females (*p* = 0.05). The presence of females with homozygosity at ANTSR suggests that ANTSR is not sufficient to trigger male development, suggesting the possibility of a second unknown CSD locus in this species. Interestingly, we found no clear pattern of increased homozygosity for diploid males near the two chromosomal homologs of *Fem* (*p* > 0.2 for both loci), suggesting that the second locus does not include a *Fem* gene, contrasting with the secondary locus in another ant species (Miyakawa and Mikheyev 2015).

**Figure 4.**
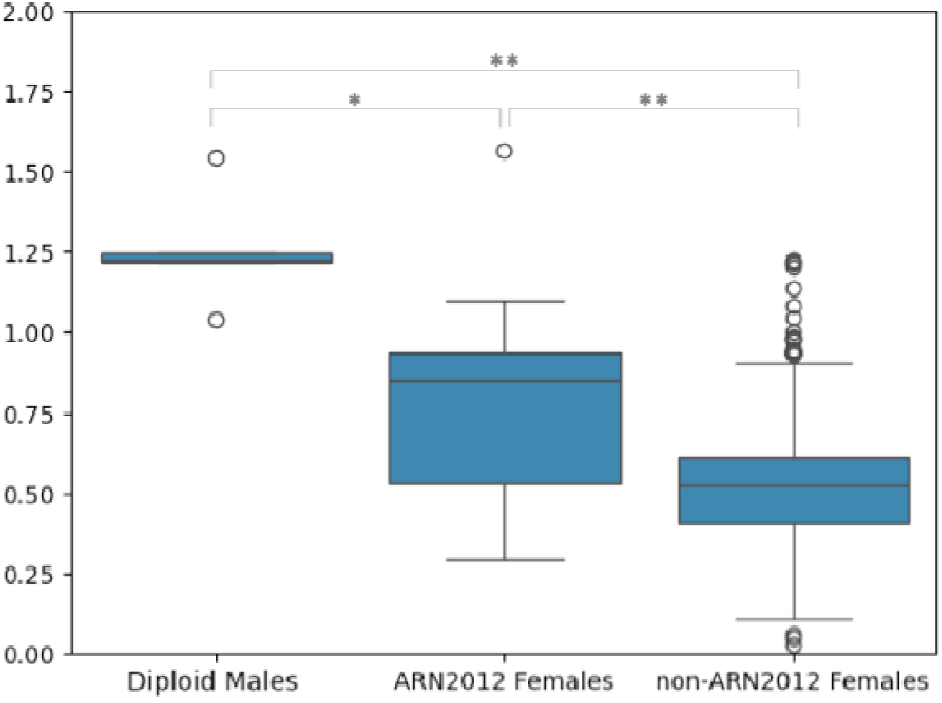
Elevated rates of homozygosity flanking ANTSR in diploid *F. cinerea* males and their female nestmates. Distance between nearest upstream and downstream heterozygous SNPs flanking the ANTSR region are shown for four males (3/4 from the ARN2102 colony), nine females from the ARN2102 colony, and 765 other females.

## DISCUSSION

Recent years have brought a flood of candidate and confident sex chromosome-linked sex determination loci, allowing for an excellent understanding of mode and tempo of turnover of sex determination in diploid organisms (Zhu et al. 2025; Kitano et al. 2024). However, whether these emerging understandings apply to haplodiploid organisms that utilize complementary sex determination (CSD) has not been clear. On one hand, some theory predicts that the rise of alternative sex determination genes in diploids may be costly (due to sex ratio skew; Bull 1983) but favored in CSD (due to reduction in production of aberrant diploid males; Crozier 1971; De Boer et al. 2008), suggesting that sex determining loci in CSD could be more fluid. On the other hand, whereas mechanistic origins of a new diploid sex determination locus can be straightforward (involving as little as gene overexpression or a novel expression pattern), origins of a new CSD locus requires mechanistic innovations in several different alleles tightly spaced in evolutionary time. Yet another possibly important set of differences involves the association of many diploid sex determination loci with long non-recombining sex chromosomal regions (e.g., Blaser et al. 2013), which does not appear to be the case under CSD; substantial data suggests that non-recombining regions accumulate costly mutations (e.g., Nguyen and Bachtrog 2021; Peona et al. 2021), which is a plausible accelerator of sex chromosome turnover in diploid organisms. The current data suggest retention of an ancestral CSD locus in a very large group of haplodiploid insects over very long evolutionary times, consistent with the perspectives that new CSD loci are hard to evolve and/or that the lack of costly non-recombining regions fails to hasten turnover under CSD, compared to diploid sex chromosomal systems.

The finding of diploid females with long runs of homozygosity surrounding the ANTSR locus in *Formica* ants is also intriguing, suggesting as it does the possibility of yet another CSD locus operating in Hymenopterans. Previous studies have shown that a *Fem* family gene serves as a CSD signal in an ant and also in a bee (Beye et al. 2003; Miyakawa and Mikheyev 2015), but the totality of evidence now suggests that these cases involve convergent evolution rather than retention of an ancient *Fem* CSD gene. That both previous cases of putatively newly-evolved CSD genes derived from *Fem* suggests that the number of genes that have the ability to evolve new CSD functions may be greatly constrained. However, our failure to find homozygosity near *Fem* genes in diploid *Formica* males suggests that a third gene family may have evolved a CSD function. Nonetheless, relying as it does on only four males from a single colony, additional studies of *Formica* are obviously highly desirable.

Limitations of data quality must be acknowledged. Assembly fragmentation, incomplete annotations, and uneven sequencing depth can constrain detection of ANTSR and complicate inference of gene loss. In addition, some polymorphisms detected within the ANTSR locus have uncertain functional consequences. On the other hand, it is of note that we were able to detect clear elevation of polymorphism despite clear limitations in our methods, relying as they do on available single non-reference isolates. Limitations notwithstanding, the complete pattern of clearly elevated polymorphism across a wide variety of Aculeata, paired with elevated polymorphism of daughters relative to their diploid brothers in *B. terrestris*, is exactly as expected by the hypothesis that ANTSR serves as the ancestral CSD locus, and difficult to explain otherwise.

This work provides an ironic coda to the most developed historical debate over the evolutionary history of CSD loci in Hymenoptera, which focused on the age of the significance of the various duplicate copies of the core insect determination gene *Fem* that are found in diverse Hymenoptera (Miyakawa and Mikheyev 2015; Koch et al. 2014). One such duplicate, *AmCSD*, was the first CSD gene to be discovered, in the honeybee *Apis mellifera*. A debate then followed as to duplicate *Fem* copies found in other Hymenoptera indicated an ancient CSD gene, with divergent perspectives seeing *AmCSD* as either very ancient or quite recently evolved (Miyakawa and Mikheyev 2015; Koch et al. 2014). The current work is consistent with the notion of an ancient CSD locus, though not with that locus being a *Fem* duplicate. Further results should interrogate the turnover of ANTSR in various Aculeata, CSD loci outside of Aculeata, and the molecular mechanisms of ANTSR’s peculiar requirement of heterozygosity for gene expression.

## METHODS

### Synteny analysis

To evaluate conservation of the ANTSR genomic region across Hymenoptera, we surveyed publicly available whole-genome assemblies and corresponding annotation files obtained from the NCBI database. In total, 112 genome assemblies spanning major Hymenopteran lineages were included in the analysis. Following Pan et al. (2024) eight protein-coding genes were assessed: GOLM1, HCF, COPA, CRELD2, THUMPD3, RPF1, BET1 and KANK3.

Available genomes varied substantially in assembly quality and annotation completeness, necessitating different approaches for assessing synteny. For species with annotated genomes, synteny was assessed by examining the order and orientation of annotated genes flanking the ANTSR locus. A species was scored as syntenic when all 8 assessed flanking genes were present in the expected order and orientation relative to ANTSR within the same genomic region.

For species lacking gene annotations, synteny was inferred using BLAST searches. Sequences of ANTSR-flanking genes identified in annotated genomes were used as queries to identify putative orthologs in unannotated assemblies. Coordinates of the resulting hits were examined to determine whether flanking genes occurred in close proximity and in a conserved arrangement. When flanking genes were located on separate contigs or assembly fragmentation prevented reliable inference, the species was classified as unresolved rather than lacking synteny. Using these criteria, syntenic conservation of the ANTSR locus was confirmed in 63 species.

### Nucleotide diversity analysis

To assess patterns of molecular diversity at the ANTSR locus, whole-genome resequencing data were retrieved from the NCBI Sequence Read Archive for species with confirmed synteny at the ANTSR locus. For each species, sequencing reads were mapped to the corresponding reference genome assembly using Bowtie. Available resequencing efforts varied in their source (single male or female, or pooled individuals) as well as the quality of their annotation (in some cases ambiguous sex or source was found). Therefore, rather than calling SNPs (which may miss variation in pooled data), we used an approach that was agnostic to the expected allelic frequency structure of the re-sequencing data. After first excluding dubious sites (sites with local indels, abnormally low- or high-coverage, or more than two nucleotides observed), we calculated estimated divergence at each site as simply the fraction of reads that had an alternative nucleotide, and this average divergence was plotted in 10kb windows.

For species with confirmed synteny, nucleotide diversity within the ANTSR region was visually compared directly to background variation in adjacent regions defined by the conserved flanking genes CRELD2 and THUMPD3. This comparative approach enabled identification of localized peaks of nucleotide diversity within or just flanking the intergenic region. Species showing clear, localized elevations in diversity were classified as exhibiting elevated polymorphism, while species with weaker or noisy signals due to limited sequencing depth, uneven coverage, or assembly fragmentation were classified as inconclusive or likely negative.

### Elevated diversity at the ANTSR locus in diverse Hymenoptera

To visualize patterns of molecular diversity, we plotted nucleotide diversity across genomic intervals spanning the ANTSR locus and the conserved flanking genes CRELD2 and THUMPD3 for representative Aculeate species (Figure X). In species classified as showing strong evidence, nucleotide diversity exhibited a pronounced local peak centered on the ANTSR region, with substantially lower diversity in adjacent flanking regions. This pattern indicates localized elevation of polymorphism rather than genome-wide increases in variation.

### Comparison of diploid males and females in B. terrestris

Three colonies of *Bombus terrestris* were supplied by the comercial breeder Biobest N.V. and maintained under controlled laboratory conditions (28/30 °C; 60 % relative humidity) in a climate-controlled room at the Zoology laboratory of the University of Mons (Belgium; Gérard et al. 2014). Among these colonies, only one produced first-generation sexuals, yielding 11 queens and 21 males. Newly emerged sexuals from this colony were transferred to in a flight cage to promote sibling mating. Successfully mated queens were subsequently used to establish second-generation inbred colonies. Over a two-month period, queens were overwintered following the protocol described by Lhomme et al. (2013). After hibernation, queens were reactivated using CO□ narcosis (Roseler 1985) and placed individually in small rearing boxes containing two workers and a cocoon in order to enhance colony initiation success(Kwon et al. 2003).

Male ploidy was determined by flow cytometry using a PA-I flow cytometer (PARTEC©; Partec GmbH, Munster, Germany). Specimens were euthanized by deep freezing at – 80°C, after which heads were dissected and immersed individually in 1 ml of DAPI-containing DNA staining solution. Sample preparation, cell cycle analysis and data quantification followed the methodology outlined by Cournault and Aron (2008).

For two of the colonies diploid sons and daughters were separately pooled (one leg per individual) and batch DNA-extracted and subjected to paired-end Illumina sequencing using standard methods (4 libraries total). For each library, reads were mapped to the reference genome using minimap2 with default parameters and the number of times that each A/C/G/T nucleotide was observed at each site was tabulated. Dubious sites were excluded (those with particularly low or high read depth or with three observed nucleotides) and sites with segregating polymorphism in females called based on at least 10% minor allele frequency supported by at least two reads. For each such site, minor allele frequency diploid sons was estimated. Minor allele frequencies were then averaged over 1Mb windows.

### Comparison of Formica females and diploid males

We downloaded data all from workers and males as well as gynes from ARN2102 from Scarparo et al. (2023), reads mapped to GCA_009859135.1 using bowtie2, and SNPs called using bcftools with standard settings. Nearly all males showed low genome-wide frequency of heterozygosity (<2%). Sites called as heterozygous in more than 2 males were discarded as dubious. Males were then assessed for heterozygosity at remaining SNPs, and four clear outliers identified (with >10% heterozygous sites) as diploid males. For each individual, the maximum genomic region overlapping ANTSR containing only homozygous sites was identified (i.e., from the nearest upstream heterozygous site to the nearest downstream heterozygous site) by a custom script. (These regions are analogous to ROH tracts, but upwardly biased given the sparsity of markers in RAD-seq data). Distribution of sizes of these ROH-like regions across different classes (diploid males, ARN2012 females, other females) were compared in R using a Mann-Whitney U test. *Fem* homologs were identified by a TBLASTN search of the *L. humile Fem* protein XM_012363646.1 against the *F. cinerea* genome, and ROH-like analyses conducted as for ANTSR.

ANTSR: 
CSD: Complementary Sex Determination

---

Supplemental Figure 1. Nucleotide diversity plots for all examined Aculeata species at local (≤250kb, left) and longer (≤2.5Mb, right) scales. The longer scale is not available for short scaffolds. Species and contig number are shown above each plot or pair of plots.

