## Supplemental Figure 1 for "Evidence for an ancient master sex determination gene in Hymenoptera"

*Apis cerana* NC_083853.1

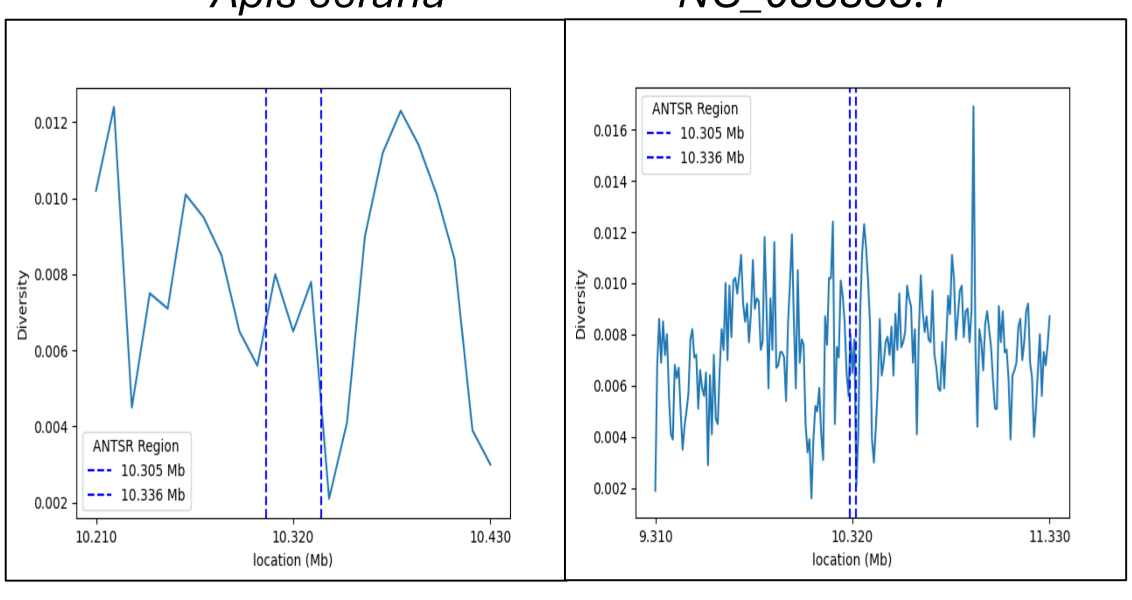

*Apis laboriosa* NW_025221207.1

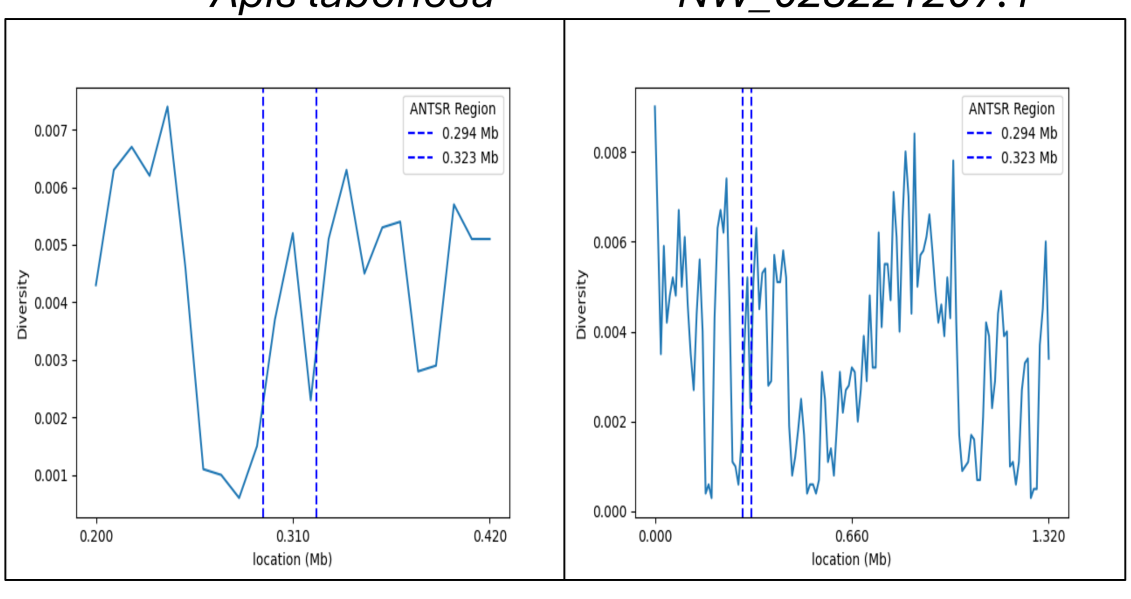

*Apis mellifera* NC_037639.1

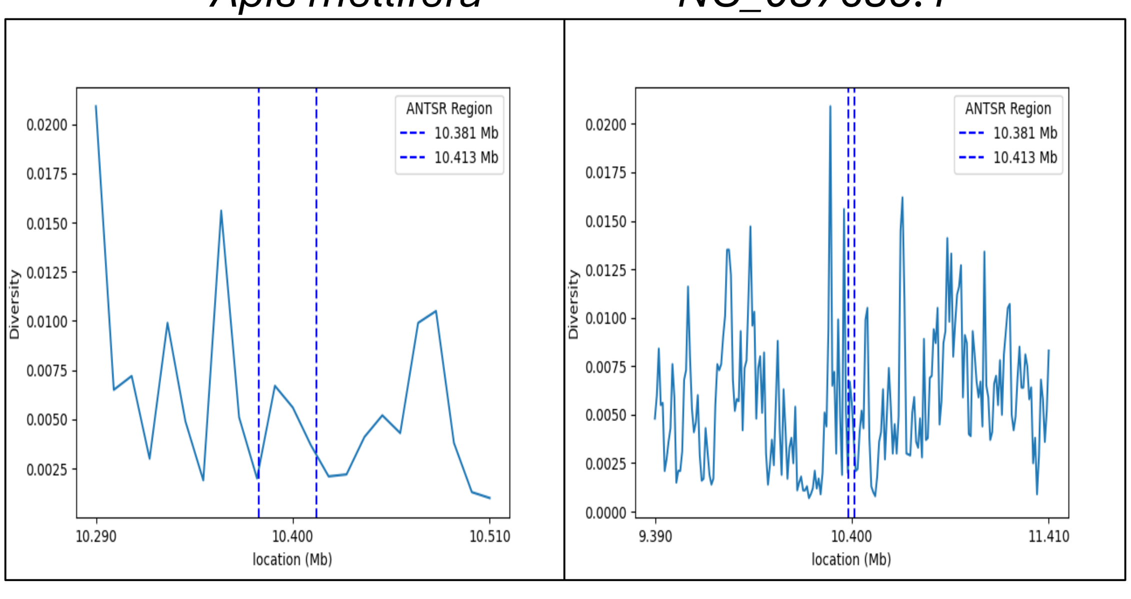

*Atta cephalotes* NW_012130075.1

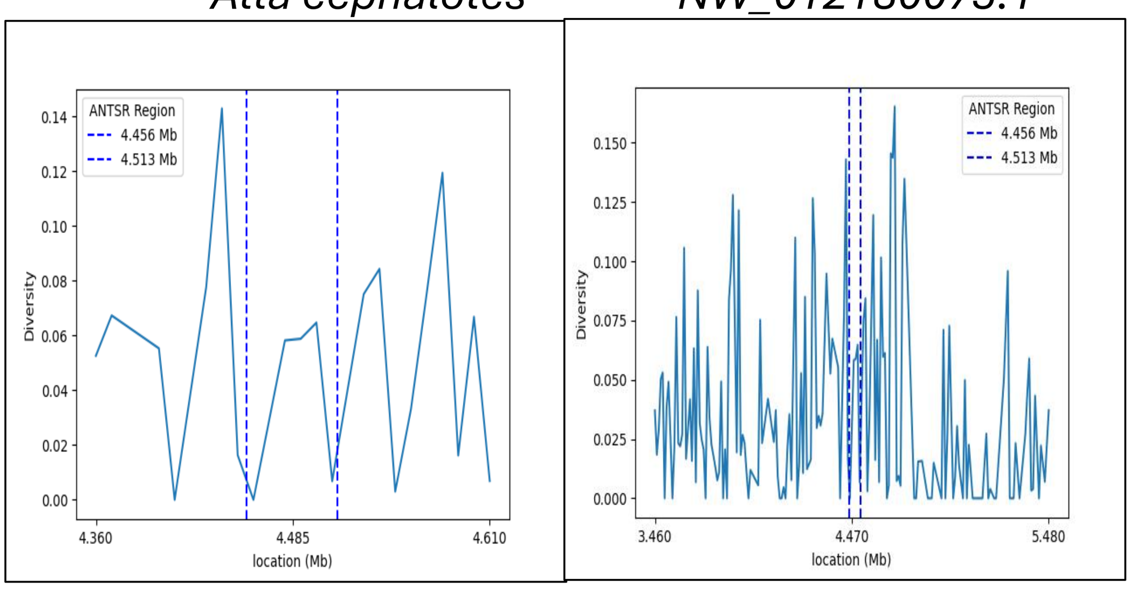

*Atta colombica* NW_017259194.1

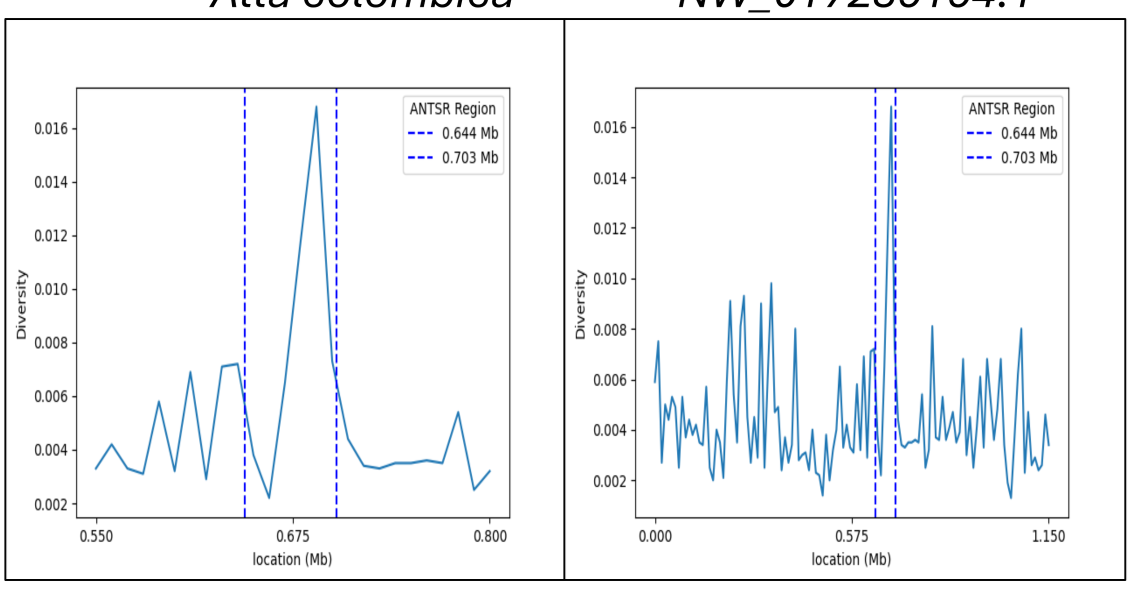

*Bombus bifarius* NW_022884386.1

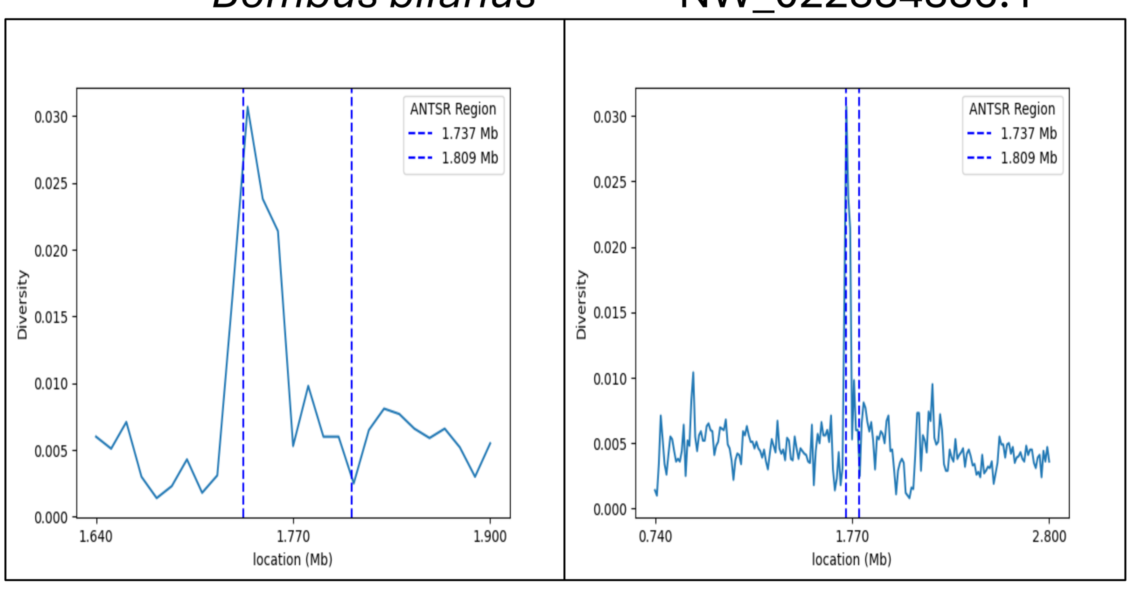

*Bombus impatiens* NT_176600.1

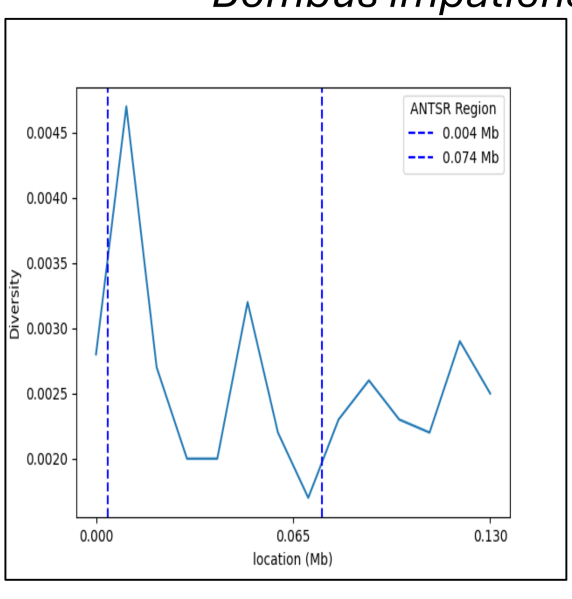

*Bombus pyrosoma* NC_057771.1

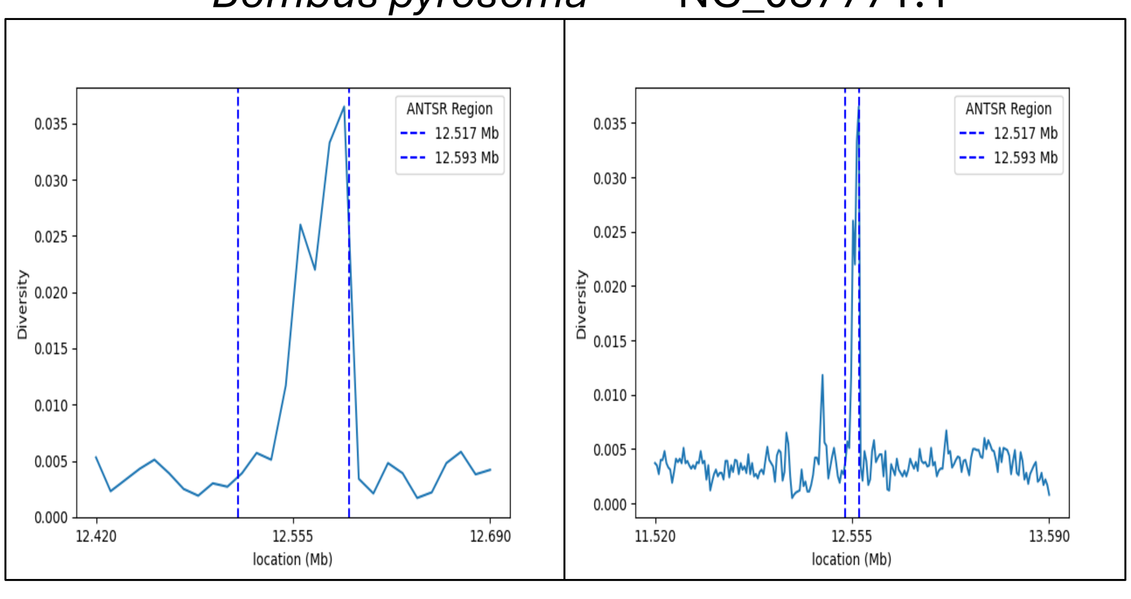

*Bombus terrestris* NC_063270.1

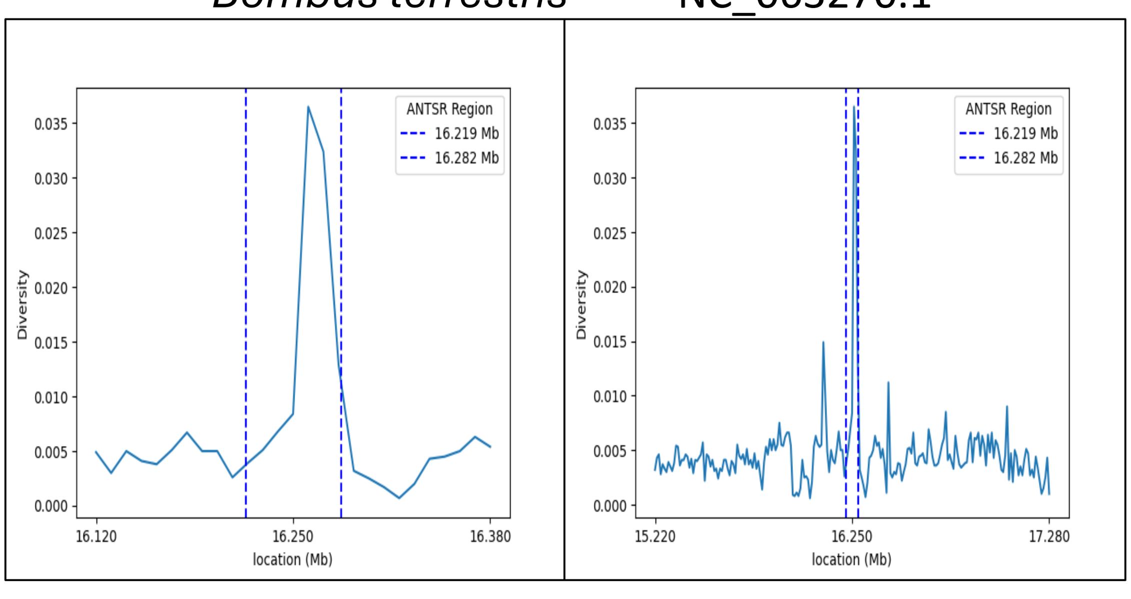

*Bombus vancouverensis nearcticus* NW_022881831.1

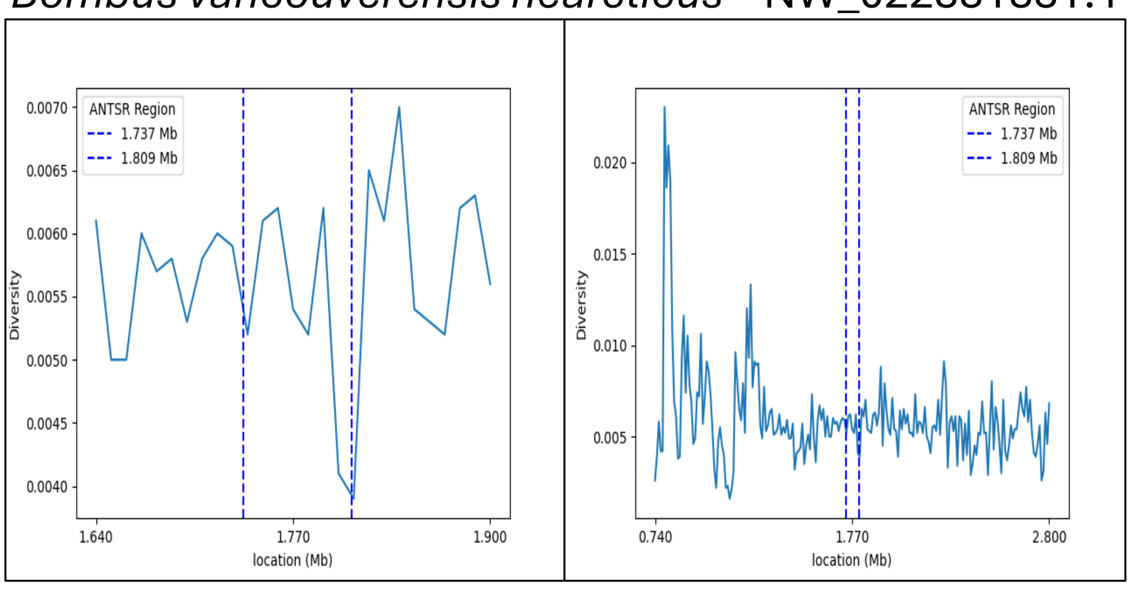

*Colletes gigas* NW_025106639.1

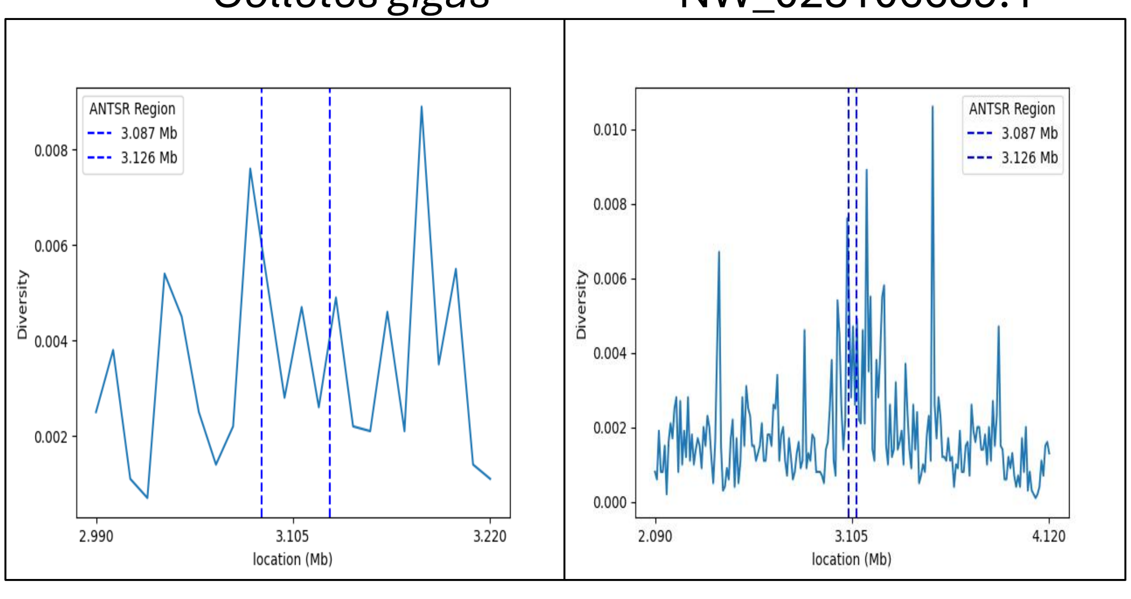

*Dinoponera quadriceps* NW_014554985.1

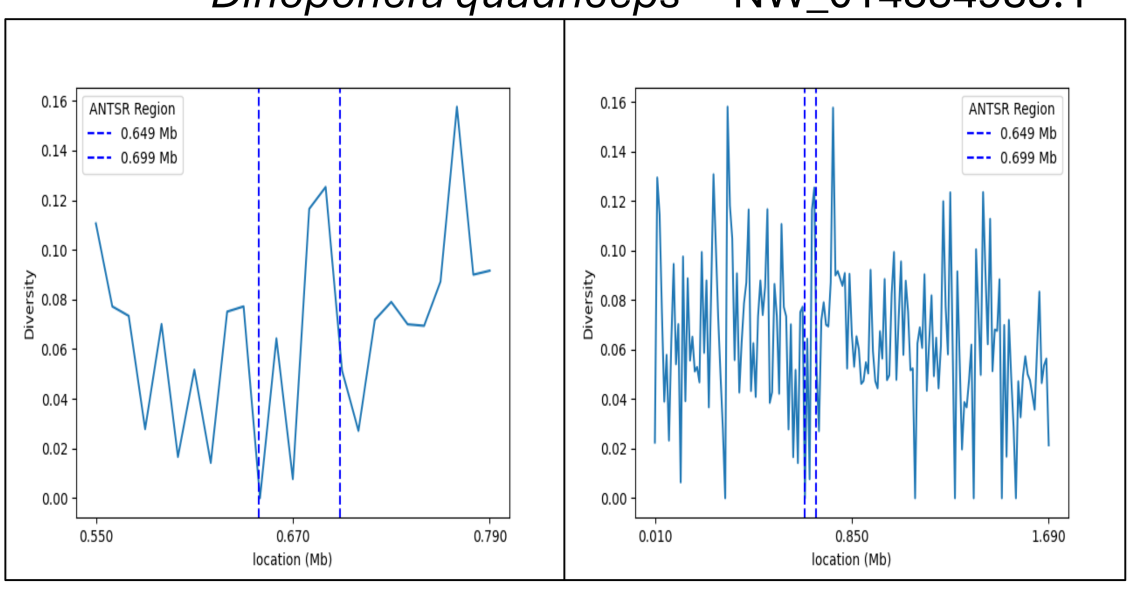

Eufriesea mexicana NW_016910897.1

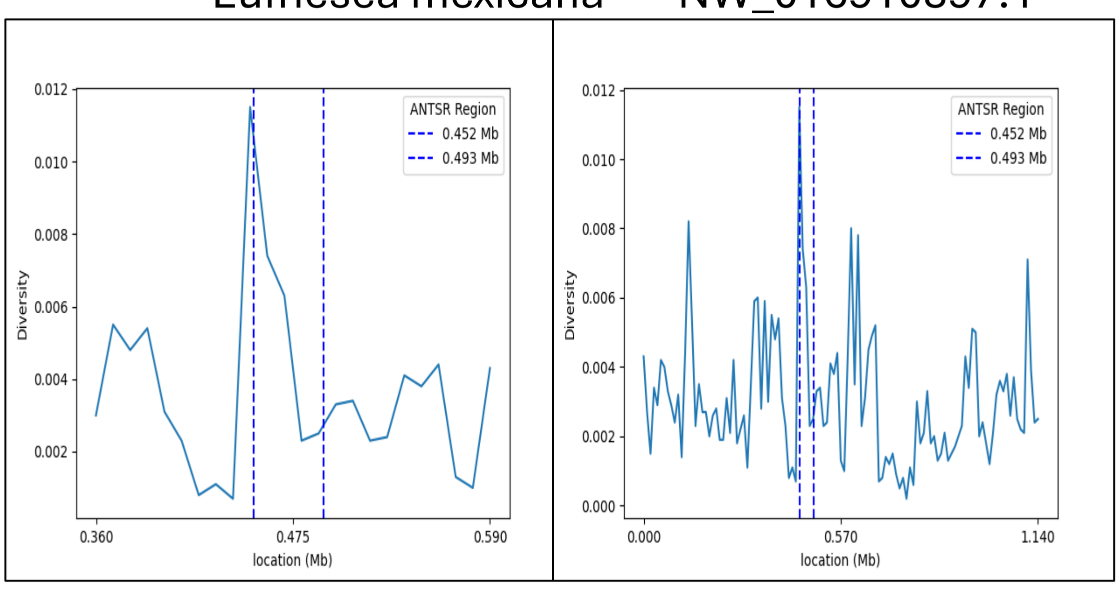

*Friesomelitta varia* NW_025110301.1

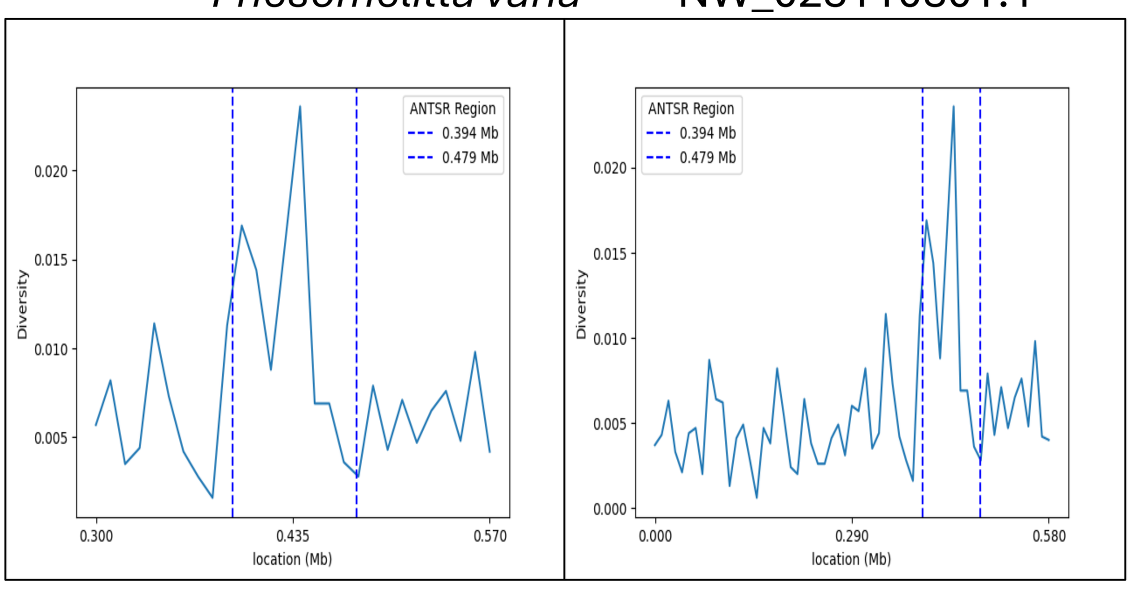

Harpegnathos saltator NW_020230332.1

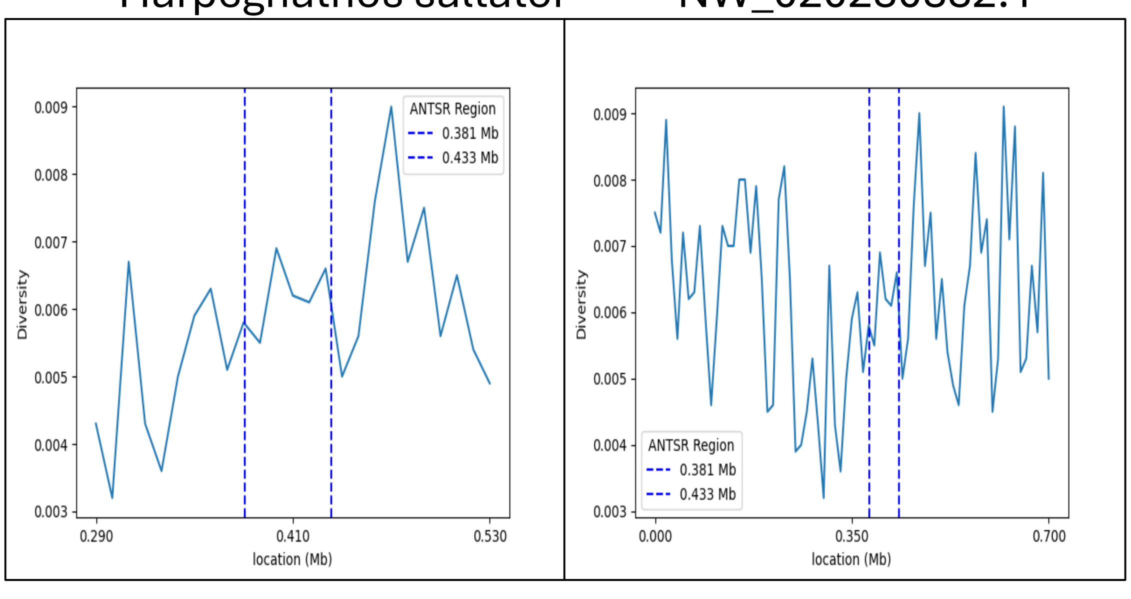

Hylaeus anthracinus NW_026534337.1

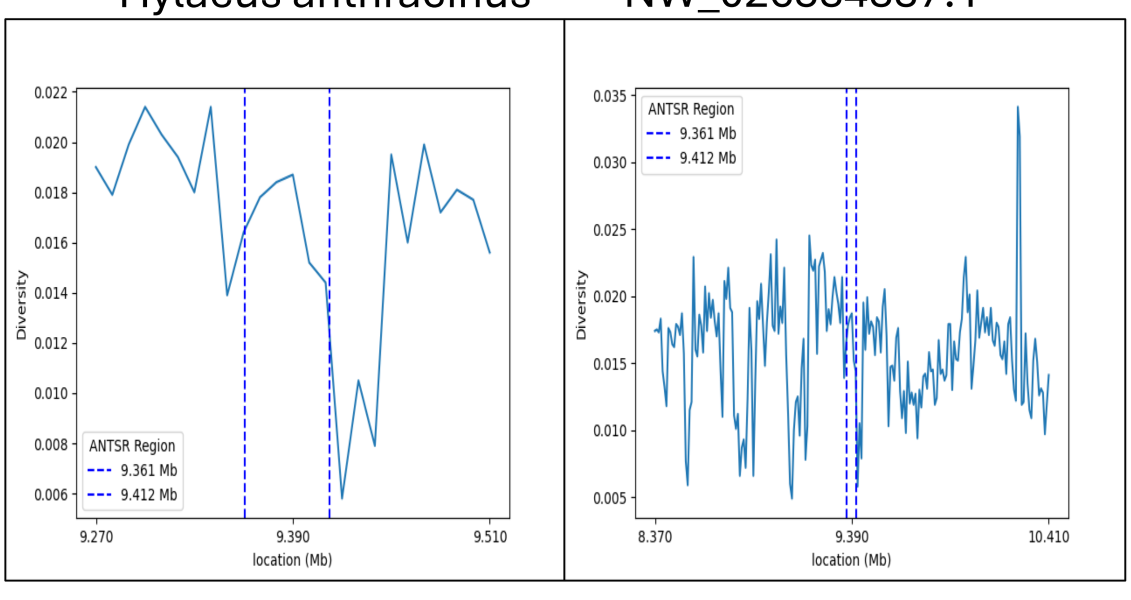

Linepithema humile NC_090131.1

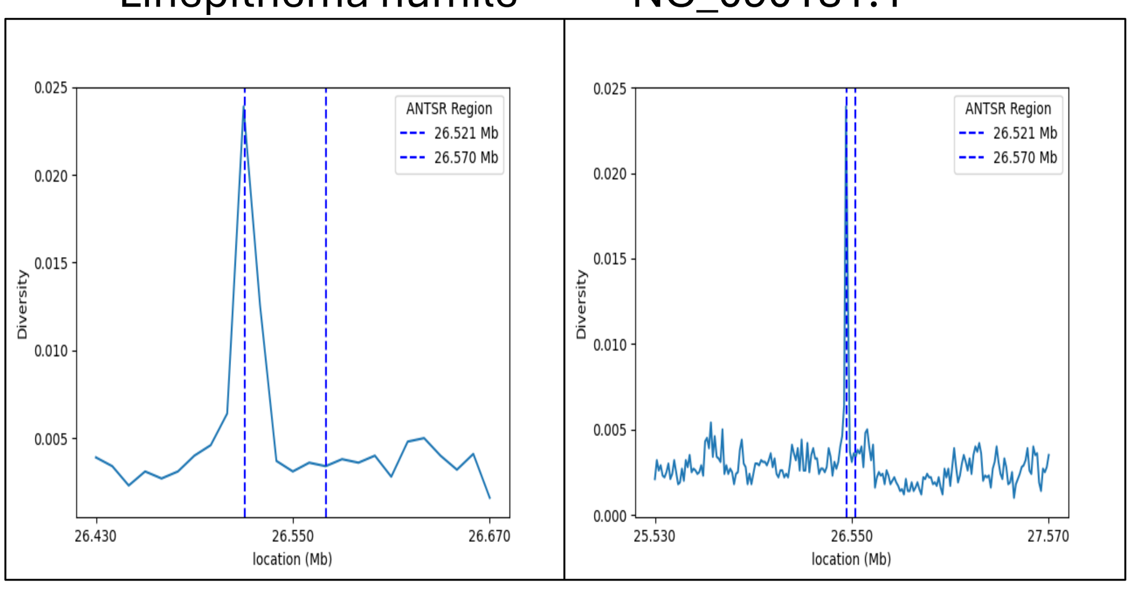

Megachile rotundata NW_003797102.1

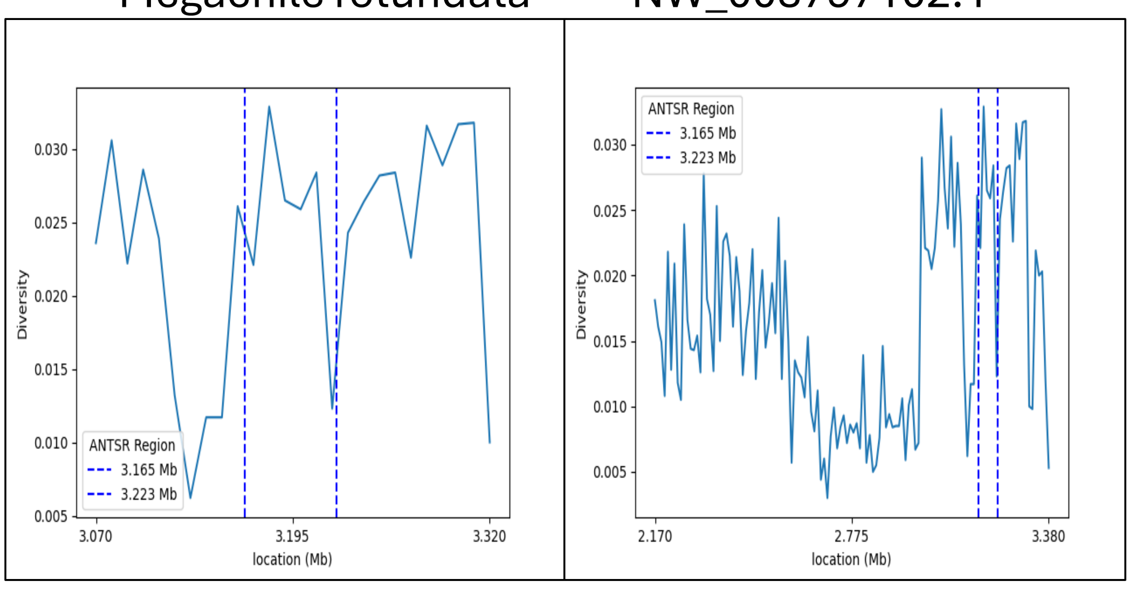

Monomorium pharaonis NC_050476.1

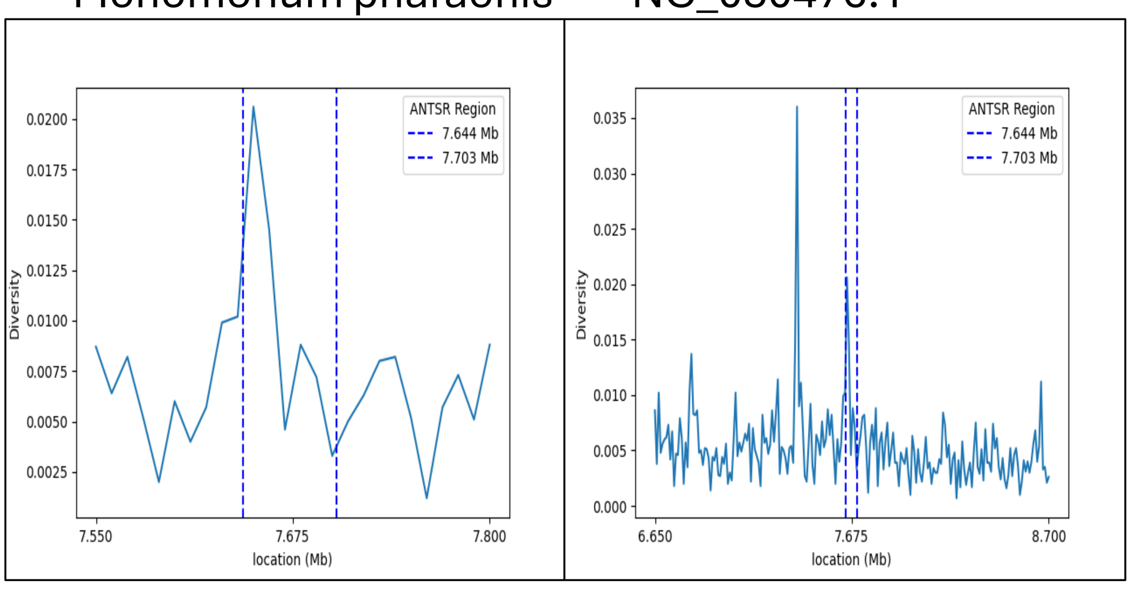

Odontomachus brunneus NW_022639499.1

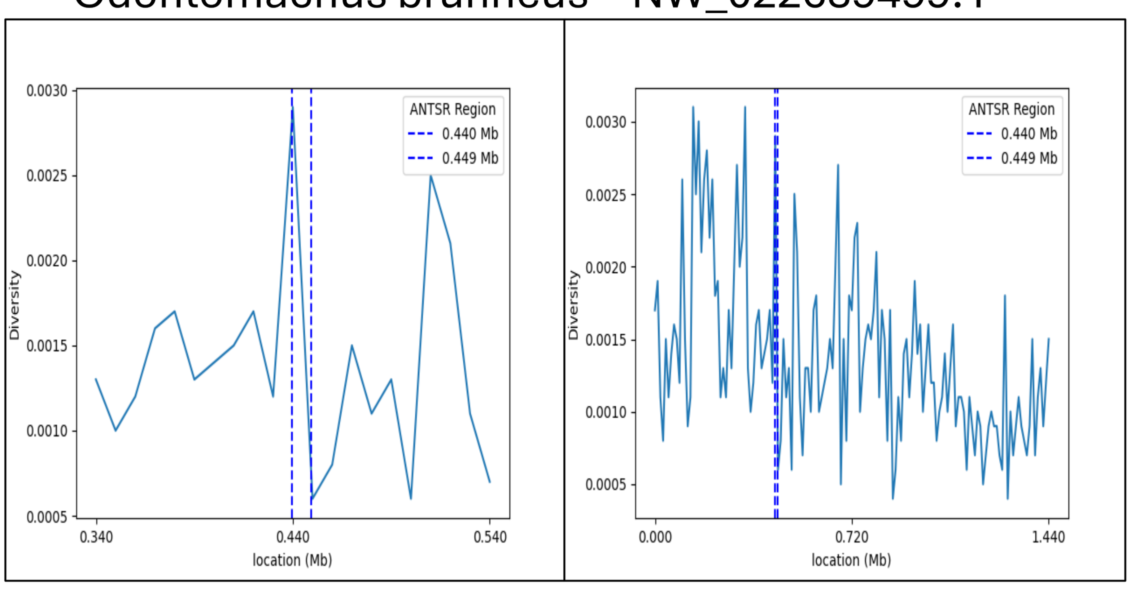

Oocerea biroi NC_039509.1

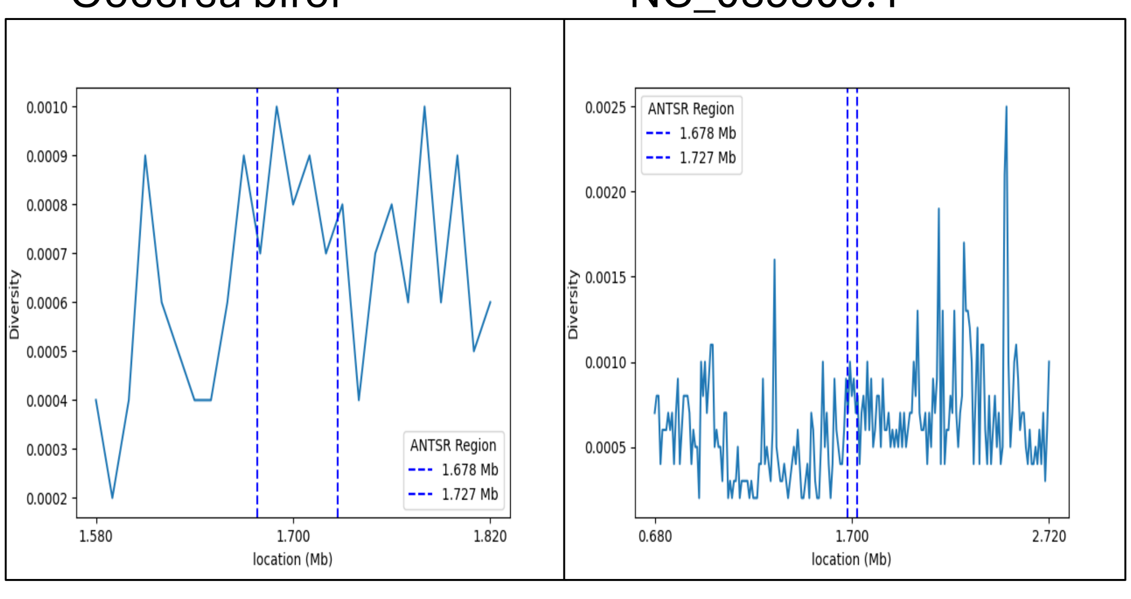

Osmia bicornis bicornis NC_060216.1

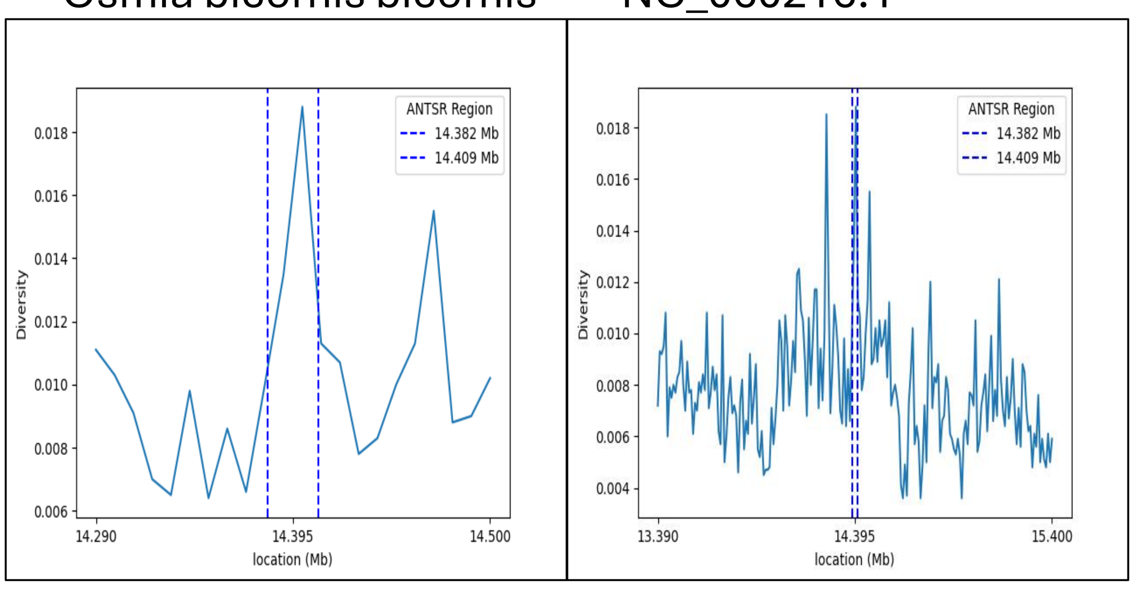

Polistes dominula NW_014569585.1

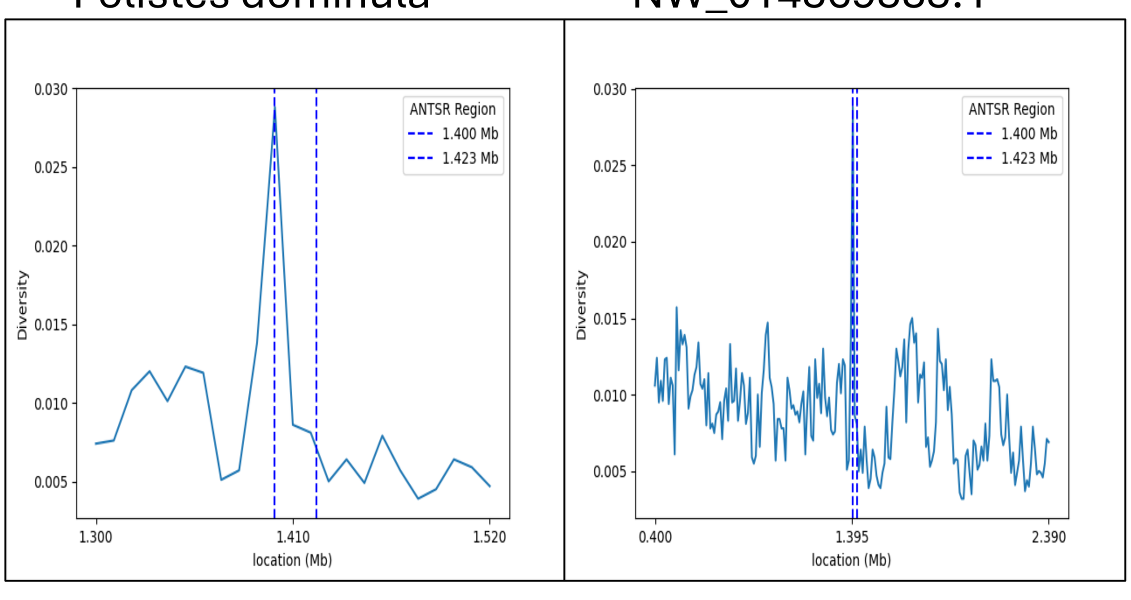

Polistes fuscatus NW_014569585.1

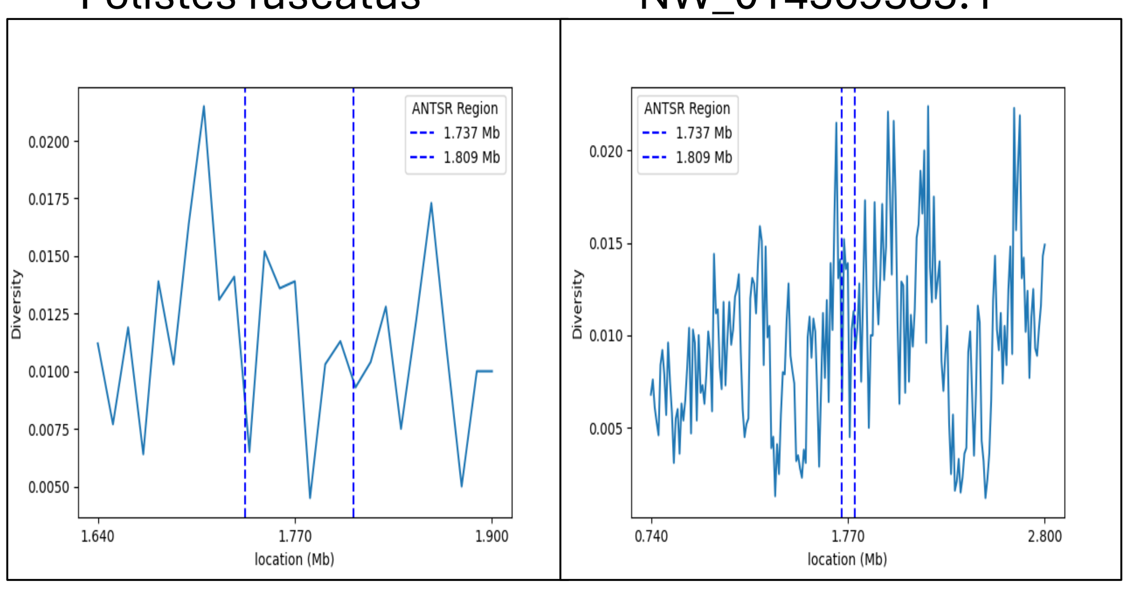

Pseudomyrmex gracilis NW_018020625.1

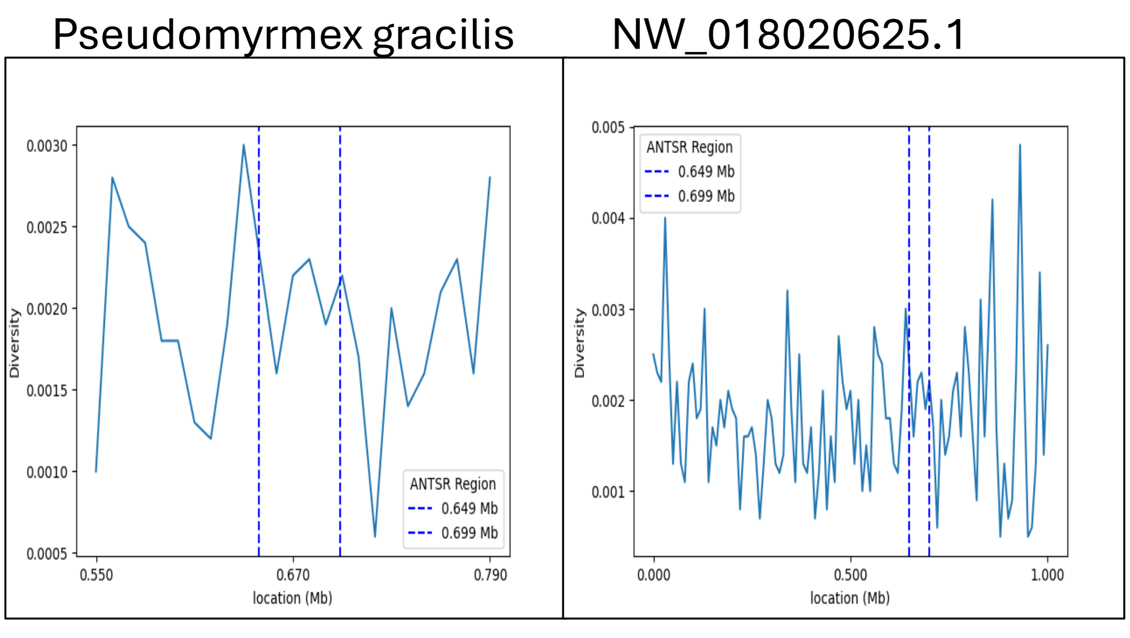

Solenopsis invicta NC_052666.1

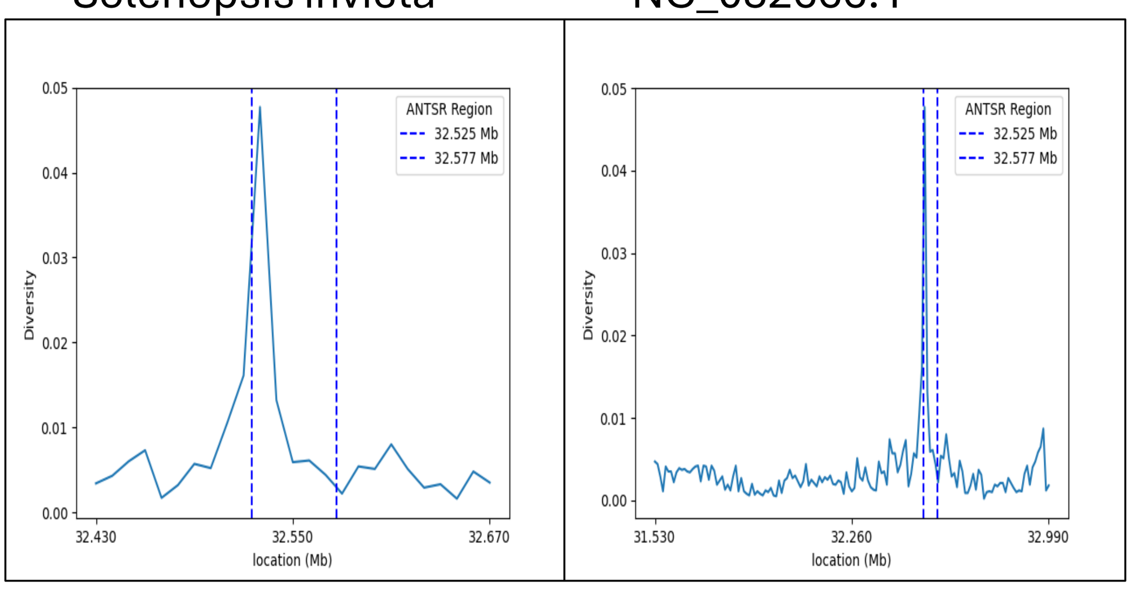

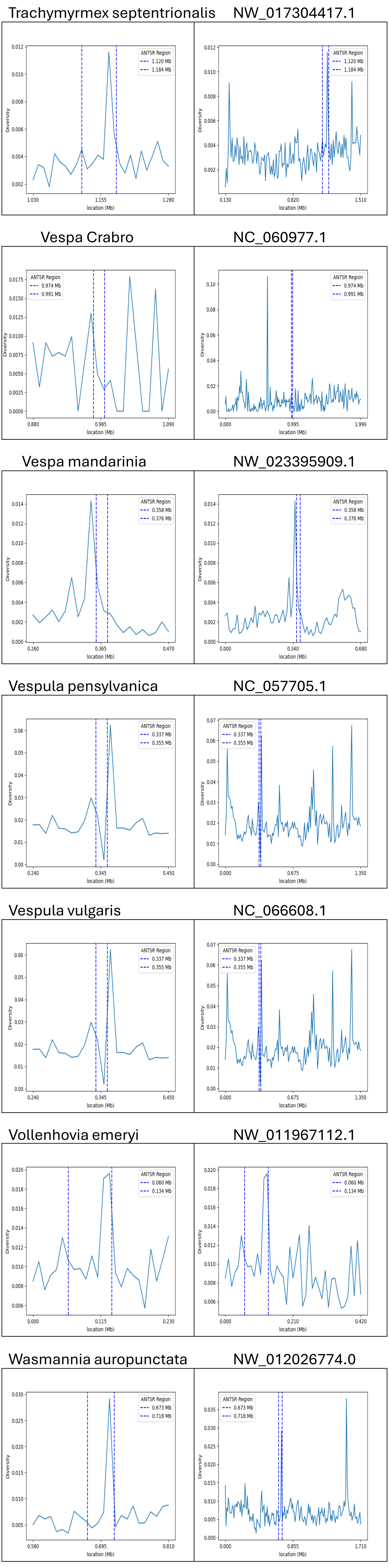
Trachymyrmex septentrionalis
NW_017304417.1
ANTSR Region
1.120 Mb
1.184 Mb
Vespa Crabro
NC_060977.1
ANTSR Region
0.974 Mb
0.991 Mb
Vespa mandarinia
NW_023395909.1
ANTSR Region
0.358 Mb
0.376 Mb
Vespula pensylvanica
NC_057705.1
ANTSR Region
0.337 Mb
0.355 Mb
Vespula vulgaris
NC_066608.1
ANTSR Region
0.337 Mb
0.355 Mb
Vollenhovia emeryi
NW_011967112.1
ANTSR Region
0.060 Mb
0.134 Mb
Wasmannia auropunctata
NW_012026774.0
ANTSR Region
0.673 Mb
0.718 Mb
Diversity
location (Mb)
